# A novel neuroelectrophysiological age index implicates brain health and sleep disorders

**DOI:** 10.1101/2022.01.24.477464

**Authors:** Soonhyun Yook, Hea Ree Park, Claire Park, Gilsoon Park, Diane C. Lim, Jinyoung Kim, Eun Yeon Joo, Hosung Kim

## Abstract

Sleep architecture and microstructures alter with aging and sleep disorder-led accelerated aging. We proposed a sleep electroencephalogram (EEG) based brain age prediction model using convolutional neural networks. We then associated the estimated brain age index (BAI) with brain structural aging features, sleep disorders and various sleep parameters. Our model also showed a higher BAI (predicted brain age minus chronological age) is associated with cortical thinning in various functional areas. We found a higher BAI for sleep disorder groups compared to healthy sleepers, as well as significant differences in the spectral pattern of EEG among different sleep disorders (lower power in slow and *ϑ* waves for sleep apnea vs. higher power in *β* and *σ* for insomnia), suggesting sleep disorder-dependent pathomechanisms of aging. Our results demonstrate that the new EEG-BAI can be a biomarker reflecting brain health in normal and various sleep disorder subjects, and may be used to assess treatment efficacy.

## 1. Introduction

Over the past decades, research has revealed the dynamic neuroelectrophysiological changes during sleep within a cyclic alternating pattern of different sleep stages.^1–3^ Stage 1, 2, and 3 are collectively known as non-rapid eye movement (NREM) sleep, which were distinguished by the presence of sleep spindles, K-complexes, and high amplitude delta waves. Rapid eye movement (REM) sleep is characterized by fast “desynchronized” brain waves with low to absent muscle tone.^2^ These typical sleep architecture and microstructures are altering with normal aging.^4–8^ Specifically, with increasing age, the percentages of stage 2, 3 and REM sleep decrease while that of stage 1 sleep increases.^2^ Previous studies have also observed decreases of sleep spindles^8,9^ and less phase coupling between sleep spindles and slow oscillations at older age.^8^ These age-related changes during different sleep stages and cycles are too complex to be estimated using a linear model.

The pattern of brain waves on sleep-electroencephalogram (EEG) in individuals with sleep disorders deviates from “normal” aging sleepers.^10–12^ Obstructive sleep apnea (OSA) is one of the most common sleep disorders and characterized by recurrent upper airway obstruction during sleep.^13–16^ The main pathology of OSA includes chronic intermittent hypoxia during sleep and sleep fragmentation that result in destructive change in sleep^13,15–18^ and increases in oxidative stress and a chronic inflammatory state.^15,17,19,20^ Insomnia is another common sleep disorder, defined by difficulty in initiating or maintaining sleep,^17^ which is characterized by a slow wave activity (SWA) deficiency as well as a hyperarousal of the central nervous system at night.^11,17,21^ By imposing hypoxic burdens and altered electrophysiological activity on the brain, these two sleep disorders may accelerate brain aging.

Machine learning techniques have been developed recently to estimate brain age index (BAI), determined by subtracting chronological age from brain age predicted from neurobiological data.^8^ The BAI has emerged as a powerful biomarker of an individual’s brain health by measuring the extent from which an individual deviates from healthy, “normal” aging trajectories.^22^ Magnetic resonance imaging (MRI) has been a popular modality to predict brain “structural” aging,^8,22–24^ but may not be sensitive to aging in the context of sleep. On the other hand, sleep EEG is more accessible to patients with sleep disorders, and reflects sleep physiology and the related brain function.^8^ Recently, estimating individual BAIs using sleep EEG have been proposed, and these approaches successfully predicted an individual’s risk for neurodegenerative disease, psychiatric disease, cognitive impairment, and mortality.^8,25,26^ Yet, an *unbiased, data-driven* deep learning (DL) approach has not been explored, which may be fully capable of modeling the complex nature of sleep neuroeletrophysiology.

Here, we proposed a DL approach to estimate individual BAIs from sleep EEG and explored the capability of this EEG-BAI as a biomarker of sleep abnormalities by associating the BAI with OSA, insomnia and comorbid insomnia and OSA (COMISA) as well as sleep parameters. We then assessed whether high EEG-BAI is associated with cortical thinning, a brain structural aging marker. As different sleep disorders may lead to brain aging through different mechanisms, we investigated their brain regional patterns of EEG spectral power.

## 2. Results

### 2.1. Accuracy of EEG-BAI model in healthy sleepers (Figure 1A)

For healthy sleepers, we hypothesized that the brain age estimated using the proposed DL model equates to their chronological age. Using a 10-fold cross validation, our DL model that predicted ages in healthy subjects (n=1,186) achieved a Pearson’s correlation coefficient of *r*=0.8 and the mean absolute error (MAE) of 5.4 years in comparison to chronological age (Figure S1). For the following analysis, we computed BAI by subtracting chronological age from predicted brain age (more details in Methods 4.3).

**Figure 1.**
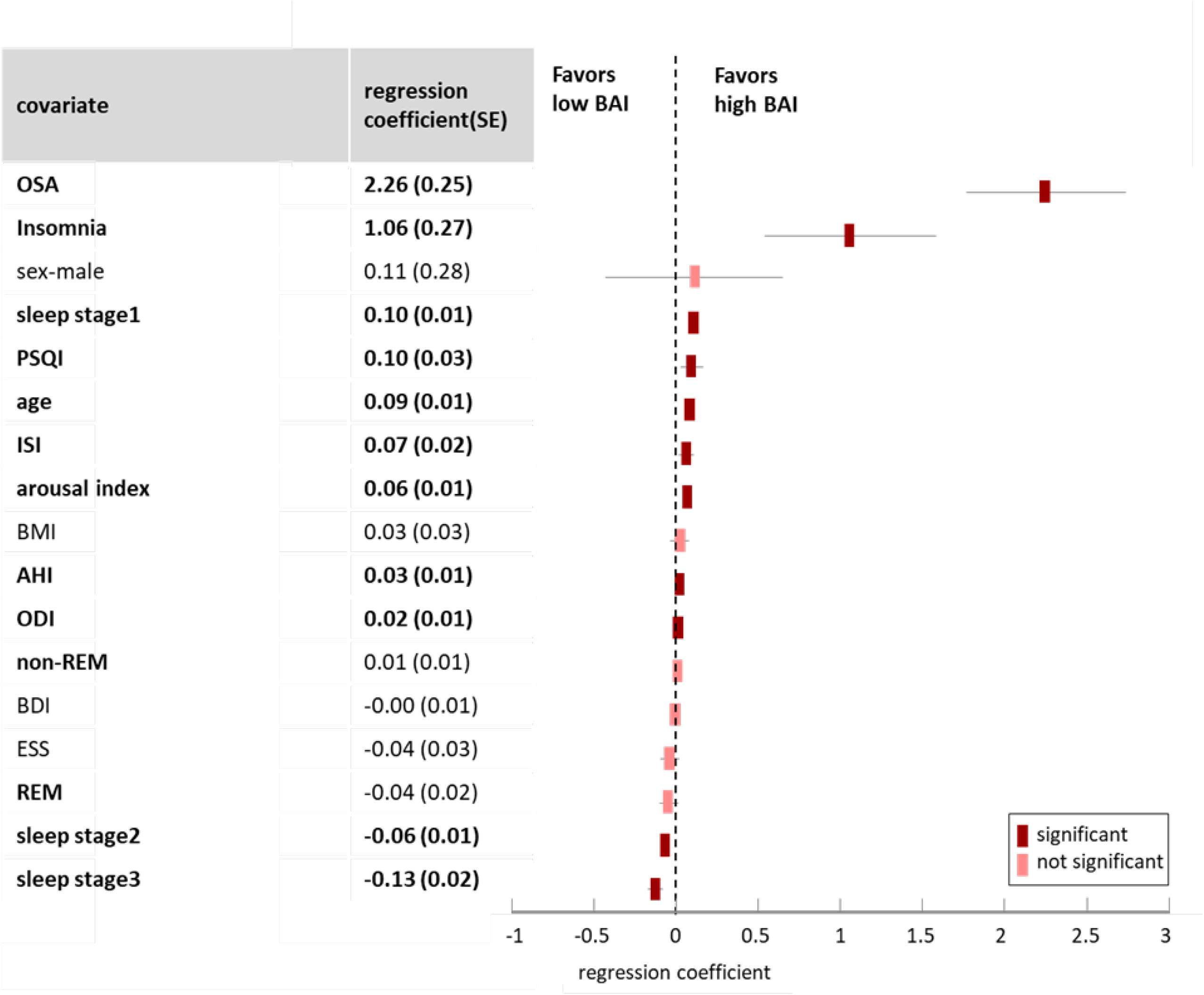
Overall scheme of brain age prediction and association between high BAI and cortical thinning. **A**. Six-channel sleep EEG data are preprocessed to remove head motion and arousal-related artifacts. Then they are converted into scalograms based on the continuous wavelet transform, resulting in 6 images in form of 16 × 2160 pixels. To build the brain age prediction model, we employed DenseNet, a type of convolutional neural network (CNN) architecture using dense connectivity. The input to the DenseNet is 6 scalogram images and the output in training was a chronological age value per individual. The model training in the current study was performed using 1,182 healthy sleepers. Mean absolute error (MAE) between chronological age and the predicted age was used as the loss function. To train and validate the brain age prediction model, a 10-fold cross validation was used. **B Top left**. Cortical thickness was calculated from the Euclidean distance between the inner surface vertices and the corresponding outer surface vertices. **Bottom left**. We parcellated the cortical surface into seven regions of interest based on human functional networks: sensorimotor, frontoparietal (FPN), dorsal and ventral attention, default mode, salient, language and auditory, and visual network. We then compared a high BAI group (n=14) and a low BAI group (n=35) that were split based on one standard deviation of BAI for the control group (7 years) to analyze the association between BAI and cortical thickness in each brain functional region. The bottom and top edges of the box indicate the 25th and 75th percentiles and the horizonal line from each square indicates 95% confidence interval. Cortical thinning for the high BAI group compared to the low BAI group was found in all different brain functional regions except for the visual network cortex. **Right**. The T-values corresponding to the brain function regions used in this group comparison are mapped on the cortical surface atlas.

### 2.2. High EEG-BAI associated with brain structural aging measured on MRI

Regional cortical thinning has been often associated with sleep disorders including OSA^16,27^ and insomnia^28^ and is also a known morphological characteristic that correlates with aging.^29–31^ To analyze the association between BAI (neuroelectrophysiological age index) and cortical thickness (brain structural age index) in various brain functional regions, we thus analyzed a sub-cohort (n=49) who underwent brain MRI, split it into two groups (high BAI vs. low BAI) based on one standard deviation of BAI for the control group (7 years) and compared cortical thickness between the high BAI group (n=14) and low BAI group (n=35). These two BAI groups had no difference in demographic and clinical characteristics (p>0.2, Table S1).

Cortical thinning was found in all the brain regions except for the visual network cortex for the high BAI group (BAI >7, determined as 1 standard deviation (SD) of the healthy sleeper group; n=14) compared to the low BAI group (BAI ≤7 years; n=35; Figure 1B). The largest cortical atrophy was observed in the language and auditory network region (*t*=2.67; *p*<.05) followed by dorsal- and ventral attention (*t*=2.61; *p*<.05), salient (*t*=2.55; *p*<.05), default mode (*t*=2.5; *p*<.05), frontoparietal (*t*=2.34; *p*<.05), and sensorimotor network regions (*t*=2.16; *p*<.05). When using a lower cut-off value (BAI=3.5) and performing the same analysis, we did not find a significant difference in cortical thickness between high and low BAI groups (*p*>0.1).

### 2.3. Higher BAI in various sleep disorder groups than healthy sleepers

Mean BAIs for OSA (mean±SD: 2.9±8.4 years), COMISA (3.3±8.5) and insomnia group (2.1±7.9) were significantly higher than healthy sleepers (*p*<0.0001; OSA, *t*=10.3; COMISA, *t*=9.4; insomnia, *t*=6.3; Figure 2A left). We also found a higher mean BAI in OSA than in insomnia and a higher mean BAI in COMISA than insomnia (*p*<0.005).

**Figure 2.**
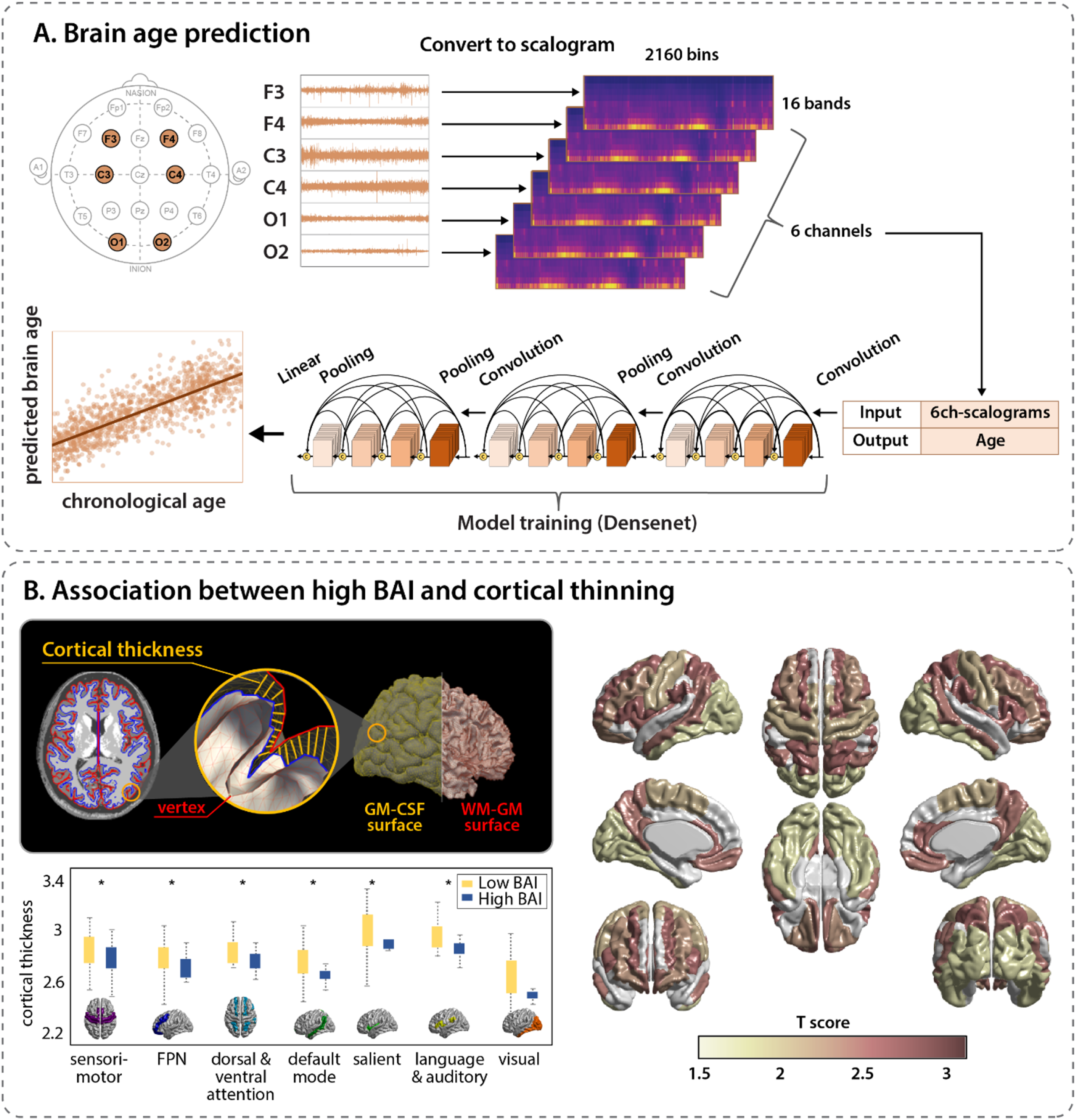
BAI of healthy sleeper vs. sleep disorder groups. **A Left**. Significantly higher BAIs were observed in all the three sleep disorder groups compared to healthy sleepers. Also, the OSA and COMISA groups displayed higher BAIs compared to the insomnia group. **Right**. The OSA, COMISA, and insomnia group displayed their own distinctive spectral patterns with respect to five frequency bands and three regions (◎: significantly different compared to the healthy sleeper group). The insomnia group presented significantly increased beta power in the frontocentral region likely due to cortical hyperarousal whereas the OSA group showed significantly decreased slower frequency bands (slow and theta) in the posterior regions, likely explaining sleep apnea-driven hypoxic burdens on cortical neural activity. Notably, the group of COMISA, a syndrome characterized by high AHI and high ISI, showed a mixed pattern of the spectral power observed in OSA and insomnia patients. **B**. Significantly accelerated brain aging over time was found in all sleep disorder groups compared to healthy sleepers. The gray solid line and gray dot line indicate mean and the 95% confidence interval of healthy sleeper, respectively. Blue, green, and yellow represent OSA, COMISA, and insomnia group, respectively. Regional spectral patterns among patient groups in comparison with healthy sleepers were computed at different ages of 30, 40, 50 and 60 years. It is clear that the sleep disorder-specific spectral patterns become more manifest as patients become older.

The spectral pattern of different wave bands for OSA, COMISA, and insomnia groups, in comparison with the healthy sleeper group, are shown in Figure 2A right. Compared to the healthy sleeper group, the OSA group presented a pattern with decreased EEG power in the waves with a frequency of 8 Hz or less; notably, the power of the slower frequency bands decreased significantly in the occipital (0.5-8 Hz) and central regions (4-8 Hz). In the insomnia group, EEG power was significantly increased compared to that of the healthy sleeper group at a frequency of 8 Hz or higher in the frontal and central regions and 12 Hz or higher in the occipital region. On the other hand, the COMISA group simultaneously exhibited characteristics of both OSA and insomnia groups. Interestingly, it revealed a significant increase of more than 15-30 Hz in the frontal and central regions, and a decrease of the 4-8 Hz band in the occipital region.

Mean BAIs for OSA and insomnia were significantly higher as chronological age increased, compared to BAI for healthy sleepers which stayed about the same at zero (Figure 2B; interaction term: age × group; p<0.0001). Using different age windows (details in 4.4.2), we identified that BAI became significantly higher in OSA relative to the healthy sleeper group since chronological age of 40 years. For COMISA and insomnia, their BAIs were significantly older than healthy subjects since 45 and 50 years (Table S2). Furthermore, the three sleep disorder groups showed different spatiotemporal patterns of the EEG spectrum (Figure 2B). We computed regional spectral patterns among the patient groups in comparison with healthy sleepers (in t-value) at four different age points of 30, 40, 50 and 60 years. This identified that the sleep disorder-specific spectral patterns seen in Figure 2A became more manifest when patients became older.

### 2.4. Incidence of sleep disorders increases in older population

As observed in Figure 3, the probability of the incidence of each disorder went higher as the chronological age increased and increased in order from the lowest BAI group Q1 to the highest BAI group Q4. Also, the Q3 and Q4 groups in OSA and COMISA exhibited a higher probability of incidence than Q1 and Q2 as chronological age increased, whereas only Q4 in insomnia displayed a higher probability of incidence.

**Figure 3.**
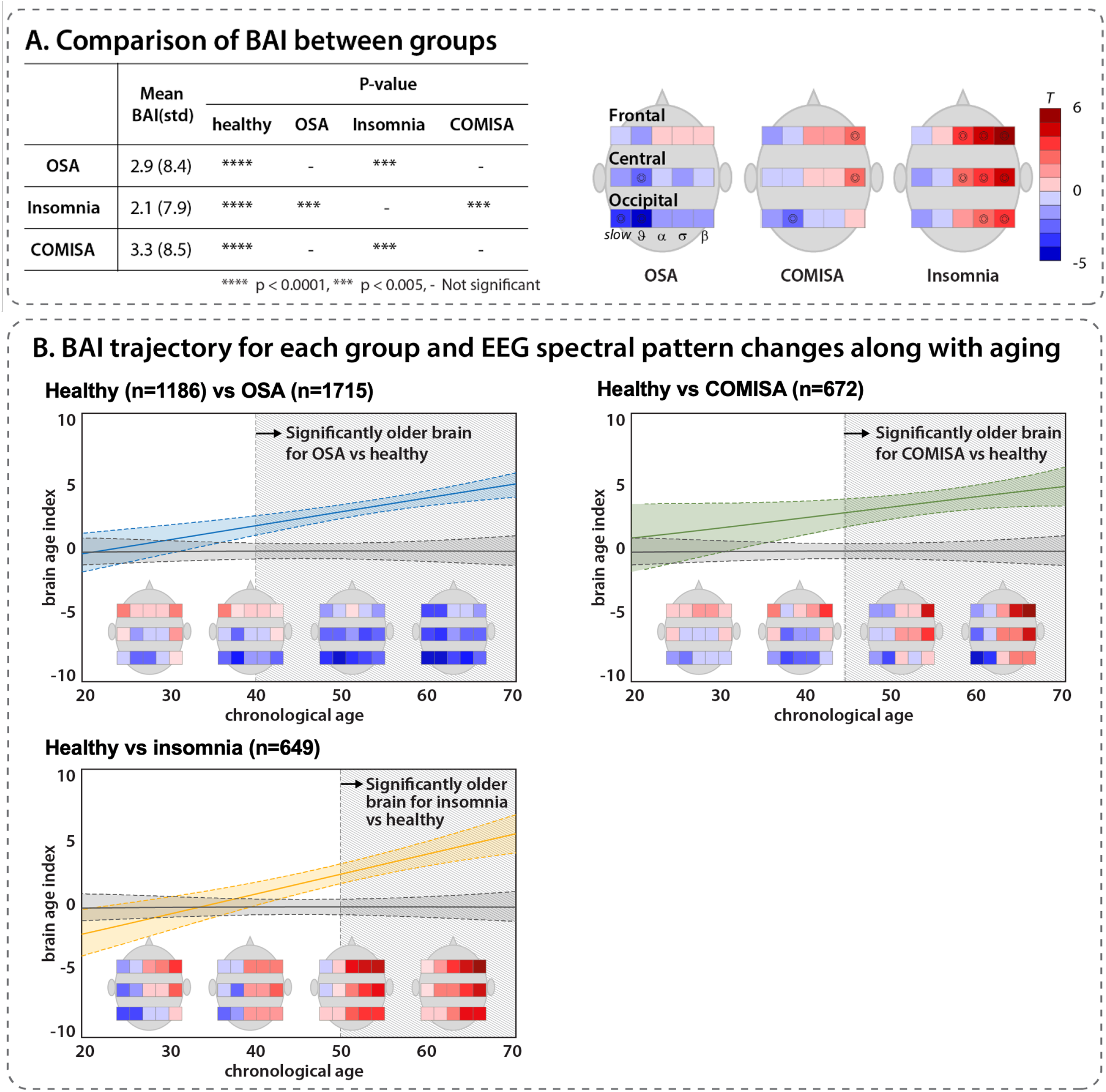
Probability of OSA & COMISA & insomnia for aging

Hazard ratios per SD for the incidence of OSA, insomnia, and COMISA were 1.0115 (CI 1.0074-1.0156), 1.0097 (CI 1.0037-1.0156), and 1.0291 (CI 1.0225-1.0357), respectively. For all three disorder groups, a higher incidence of disorder was significantly associated with a higher BAI.

### 2.5. Regression analysis of BA prediction

Coefficients of covariates in our linear regression model that estimated BAI are presented in Figure 4. The following covariates were significantly associated with a higher BAI; incidence of OSA was the highest positive regression coefficient with BAI (regression coefficient±standard error: 2.26±0.25; *p*<.001), followed by incidence of insomnia (1.06±0.27; *p*<.001), age at EEG (0.09±0.01; *p*<.001), percentage of the length of stage 1 sleep (0.10±0.01; *p*<.001), insomnia severity index (ISI) (0.07±0.02; *p*<.001), Pittsburgh Sleep Quality Index (PSQI)^32^ (0.10±0.03; *p*=.002), arousal index (0.06±0.01; *p*<.001), apnea hypopnea index (AHI) (0.03±0.01; *p*<.001), and oxgen desaturation index (ODI) (0.02±0.01; *p*<.001). The percentages of the lengths of stage 2 sleep (−0.06±0.01; *p*<.001) and stage 3 sleep (−0.13±0.02; *p*<.001) were negatively correlated with BAI. Additionally, Epworth Sleepiness Scale (ESS)^33^ was not associated with BAI, but only positively correlated for over 50 years old (1.26±0.38; *p*<.001).

**Figure 4.**
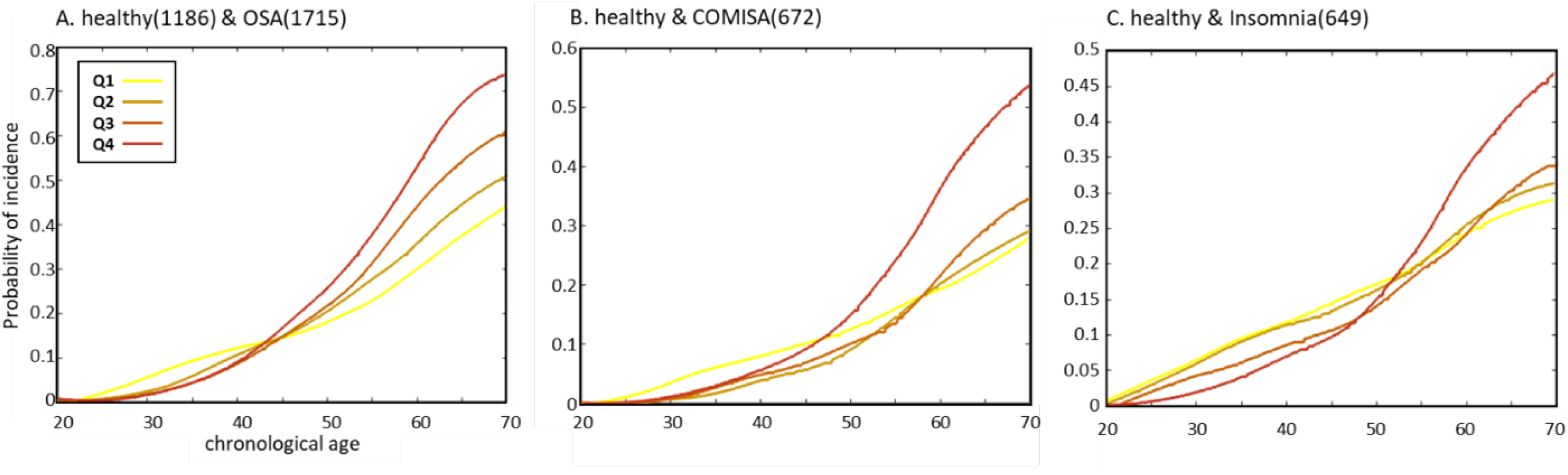
Regression analysis of BA prediction. Squares indicate coefficients of covariates and the horizonal line from each square indicates 95% CI. Abbreviations: PSQI, Pittsburgh Sleep Quality Index; ISI, Insomnia Severity Index; BMI, Body Mass Index; ODI, Oxygen Desaturation Index; BDI, Beck’s Depression Inventory; ESS, Epworth Sleepiness Scale; REM, Rapid Eye Movement

Different patterns of association between covariates and BA were observed among four groups (Table 1). The BAI of the healthy sleeper group was positively correlated with sleep stage 1 (0.19±0.03; *p*<.0001) and negatively correlated with sleep stage 3 (−0.13±0.02; *p*<.0001). In the OSA group, age (0.10±0.02; *p*<.0001) and stage 1 sleep (0.08±0.01; *p*<.0001) were positively associated whereas sleep stage 2 (−0.052±0.02; *p*<.005) and sleep stage 3 (−0.17±0.04; *P*<.0001) were negatively correlated. In the COMISA group, age (0.09±0.03; *p*<.05) and sleep stage 1 (0.07± 0.02; *p*<.005) were positively associated. In the insomnia group, age (0.14±0.02; *p*<.0001), arousal index (0.27±0.04; *p*<.0001), and sleep stage 1 (0.24±0.03; *p*<.0001) were positively associated whereas sleep stage 3 (−0.26±0.04; *p*<.005) was negatively correlated. These covariates are all significant with BAI when corrected by FDR multiple comparison.

**Table 1.** Regression Analysis of BA Covariates separately for healthy sleeper, OSA, COMISA, and insomnia

| Covariate | Healthy |  | OSA |  | COMISA |  | Insomnia |  |
| --- | --- | --- | --- | --- | --- | --- | --- | --- |
|  | Regression coefficient (SE) | *P* | Regression coefficient | *P* | Regression coefficient | *P* | Regression coefficient | *P* |
| age | 0.00 (0.01) | - | 0.10 (0.02) | *** | 0.09 (0.03) | * | 0.14 (0.02) | *** |
| BDI | -0.02 (0.03) | - | 0.01 (0.03) | - | 0.04 (0.03) | - | -0.05 (0.03) | - |
| AHI | 0.11 (0.04) | - | -0.02 (0.01) | - | -0.03 (0.01) | - | 0.10 (0.07) | - |
| ODI | 0.10 (0.05) | - | -0.03 (0.01) | * | -0.04 (0.01) | * | 0.03 (0.08) | - |
| ISI | 0.00 (0.05) | - | 0.01 (0.05) | - | 0.08 (0.11) | - | 0.09 (0.08) | - |
| Arousal index | 0.08 (0.04) | - | 0.02 (0.01) | - | 0.03 (0.02) | - | 0.27 (0.04) | *** |
| Non-REM | 0.04 (0.04) | - | 0.25 (0.02) | - | 0.00 (0.01) | - | 0.02 (0.04) | - |
| sleep stage1 | 0.19 (0.03) | *** | 0.08 (0.01) | *** | 0.07 (0.02) | ** | 0.24 (0.03) | *** |
| sleep stage2 | 0.00 (0.02) | - | -0.05 (0.02) | ** | -0.06 (0.02) | - | 0.00 (0.03) | - |
| sleep stage3 | -0.13 (0.02) | *** | -0.17 (0.04) | *** | -0.10 (0.05) | - | -0.26 (0.04) | *** |
| REM | -0.04 (0.04) | - | -0.01 (0.03) | - | -0.10 (0.04) | - | -0.02 (0.04) | - |
| sex-male | 0.80 (0.45) | - | -1.12 (0.58) | - | -0.63 (0.81) | - | -0.78 (0.62) | - |
| PSQI | -0.10 (0.07) | - | 0.06 (0.08) | - | 0.24 (0.10) | - | 0.18 (0.09) | - |

- not significant; \*\*\* $p < 0.0001$ ; \*\* $p < 0.005$ ; \* $p < 0.05$

## 3. Discussion

In this study, we proposed a sleep EEG-based brain age prediction model and demonstrated that a higher BAI was associated with brain atrophy. Faster brain aging was found in patients with sleep disorders compared to healthy sleepers, which was even more adverse in older age. We identified that various clinical factors related to sleep, including OSA, insomnia, proportions of sleep stages, PSQI, ISI, arousal index, AHI, and ODI were significantly associated with BAI. Furthermore, we found significant differences in BAI and the spatiotemporal pattern of EEG spectrum among different sleep disorders, suggesting different pathophysiology leading to unique brain aging patterns among sleep disorder groups.

### Electrophysiological brain aging vs. structural brain aging

Various brain disorders and diseases result in irreversible brain structural alterations such as decreased cortical thickness or decreased regional volume.^34–37^ We hypothesized that an EEG-BAI higher than the normal range is associated with significant brain cortical thinning on MRI. Our finding indeed demonstrated that the subgroup with an elevated EEG-BAI (BAI>7 years) displayed a significant cortical thinning compared to the group with low EEG-BAI. Interestingly, the elevated EEG-BAI group demonstrated cortical thinning in multiple areas associated with high order cognitive function (attention, language and frontoparietal) and sleep/wake networks (default mode, salient). On the other hand, when we analyzed and compared a less elevated EEG-BAI group (BAI>3.5 years) with a lower group of EEG-BAI ≤ 3.5, we observed no significant cortical thinning. This result suggests that only a sufficiently large EEG-BAI is required to drive brain structural changes. In other words, the importance of this finding lies in that brain physiological aging may reflect short-term or temporal brain health conditions. Furthermore, applying early, proper treatment to subjects with a sleep disorder to maintain a low BAI or reverse an initially high BAI may prevent brain structural degeneration and impede development of cognitive impairment. However, a longitudinal study on pre- and post-treatment effects is needed to clarify the efficacy of treatment.

### Identification of clinical factors associated with faster EEG-BAI

Regression analysis of BAI and coefficients of covariates in our study allows for clarifying the source of accelerated brain aging by determining and ranking the weight of various clinical factors associated with an increased BAI. In this analysis, having OSA and being insomniac were the two strongest factors associated with BAI increase. High PSQI and ISI values also correlated with a high BAI, supporting that sleep disturbances contribute to pathologically accelerated brain aging. Among PSG parameters, increases of apnea/hypopnea (high AHI) and oxygen desaturation (high ODI) positively correlated with a high BAI, indicating the substantial role of disordered breathing during sleep and subsequent recurrent hypoxic burden in pathologic brain aging.^15^ Furthermore, the duration of sleep stage 1 positively correlated with BAI, whereas the duration of sleep stages 2-3 and REM sleep were negatively correlated. Although a longer stage 1 sleep, shorter slow wave sleep and shorter REM sleep can occur in the normal aging process,^2^ the pathologically aggravated alterations of sleep disorder-related sleep structure may significantly reflect an atypical pattern of brain electrophysiology and be further associated with accelerated brain aging.

### Disease-specific features of sleep disorders in aspects of brain age (OSA vs. COMISA vs. insomnia)

In this study, different rates of BAI change relative to chronological age were observed among different disease groups. First, sleep disordered breathing had a greater impact on brain aging than insomnia because OSA and COMISA groups displayed a significantly greater BAI than the insomnia group, as well as significant differences in BAI compared to the healthy sleeper group from an earlier age relative to that of the insomnia group (Figure 2). This observation is possibly explained by the following: OSA and COMISA are characterized by a combination of recurrent hypoxia events as well as sleep fragmentation altering sleep structures whereas insomnia is associated only with sleep disturbance.^17^ Impaired cerebral perfusion caused by hypoxia-related endothelial dysfunction in OSA has been reported in several recent studies.^38,39^ It is plausible that the impaired cerebral perfusion could result in additional or synergistic effects on pathologic brain aging when combined with disrupted sleep structures.

EEG spectral characteristics analysis showed distinctive spectrum profiles among the three sleep disorder groups, supporting the different pathomechanisms in relation to brain aging involved in different sleep disorders. The insomnia group presented significantly increased beta power in the frontocentral region, suggesting the important role of cortical hyperarousal in pathologic brain aging in insomnia.^12^ The OSA group showed significantly decreased slower frequency bands (slow and theta) in the posterior regions, likely explaining for a poor deep sleep due to sleep fragmentation as well as a hypoxic burden on cortical neural activity due to sleep apnea.^40^ Observed as the lower spectrum of slower EEG waves, these slow frequency bands ultimately result in an increased BAI. These findings of sleep microstructural changes are consistent with previous small sample studies of OSA and insomnia patients^12,40^ in that the lower spectrum power of slower waves was observed in the OSA groups compared to controls.^41,42^ In contrast, greater alpha, sigma, and beta EEG activities in NREM^43^ and REM sleep^44^ were found in patients with chronic insomnia.

Notably, patients with COMISA, a syndrome characterized by high AHI and high ISI, showed a mixed pattern of the spectral power observed in OSA and insomnia patients. Due to worse treatment adherence patterns (e.g. adherence to continuous positive air pressure (CPAP) treatment), COMISA has increasingly become a point of interest in sleep research.^45^ Our new finding of EEG spectral patterns of COMISA, which have been rarely investigated, could be used to characterize the comorbidity of this specific disease group.

Furthermore, the disorder groups within the 20-40 years age bracket did not show any significant differences in BAI and the electrophysiological pattern (= EEG power spectrum pattern) compared to the healthy sleeper. On the other hand, in older patients, we found that the BAI in each disorder group increased with chronological age, and the sleep disorder-specific electrophysiological pattern became more manifest. It is thus compelling to suggest that faster brain aging in patients with OSA and insomnia occurs via unique disease-specific pathomechanisms.

### Limitations and future directions

The current study has some important limitations. First, our EEG data as part of the standard PSG were obtained through six sole channels; thus the spatial resolution of EEG is limited while the temporal resolution of 6 hours is high. Second, the distribution of ages in the dataset was not completely equal between healthy sleepers and patients with sleep disorders, which can potentially bias findings even if we used linear regression models to correct it. Third, a future study is required to assess the possible associations between neuroelectrophysiological aging and cognitive performance or related dementias. Fourth, our study solely contains cross-sectional data. Accordingly, we could not investigate the disease progression or effects of treatment (e.g., CPAP treatment in OSA) on reversibility of brain aging, which is important and clinically relevant. Fifth, the proposed EEG-BAI model showed lower accuracy in healthy subjects compared to image-based models (MAE: 5 vs 3 years),^22^ likely because of the nature of sleep EEG (e.g., scalp EEG is more noisy than MRI; multiple night EEGs may be required). Nonetheless, it is noteworthy that our model showed the highest performance among the existing EEG-BAI models, most of which relied on a variety of mathematically defined EEG features.^8,46^ Lastly, we did not use sleep stage information because deep learning approaches are designed to learn various information related to brain aging from raw data or least processed data. However, this process could potentially limit our interpretation as individual data were not aligned in terms of sleep stages. Recurrent neural networks such as transformers, long short-term memory (LSTM) and gated recurrent units (GRU) are possibly better to learn temporally varying data like EEG than the proposed CNN model although we incorporated a scalogram approach into the 2D CNN structure.

### Conclusion

Our study demonstrates that the neuroelectrophysiological BAI estimated using a deep learning model of sleep EEG can serve as a potential biomarker that reflects the pattern of sleep microstructure and brain health in patients with sleep disorders. Our data also suggested that disordered sleep breathing and insomnia may yield pathologic brain aging through different pathophysiological mechanisms. Thus, BAI and the related EEG spectral patterns can be used to phenotype sleep disorders as well as screen for sleep abnormalities that potentially harm brain health.

## 4. Materials and Methods

### 4.1. Subjects and EEG acquisition

Our cross-sectional dataset comprised of 5,282 (3,633 males) who underwent polysomnography at Samsung Medical Center between 2015 and 2019. These subjects did not use positive airway pressure therapy, mandibular advancement devices, or oxygen therapy during the measurement. From this dataset, we excluded subjects with a history of cerebrovascular disease, other neurological (neurodegenerative disease, brain tumor, epilepsy, or severe head trauma) and psychiatric diseases (psychosis, current depression). We also excluded a small sample of subjects who were younger than 20 years or older than 70 years as they were scattered over different ages. Each PSG is scored by one of seven EEG technicians according to the American Academy of Sleep Medicine (AASM) standards (AASM, 2007). Healthy sleeper, OSA, insomnia, and COMISA groups were defined by the insomnia severity index (ISI) as well as the apnea-hypopnea index (AHI).^47^ OSA group was defined by moderate to severe OSA (AHI ≥15) without other sleep complaints (ISI≤14), and insomnia group was defined by ISI over 14^47^ without clinically significant OSA (AHI<15). Subjects with AHI ≥15 and ISI >14 were classified into the COMISA group. Following the exclusion and classification criteria, our final study subjects included 1,186 healthy sleepers (751 males; median ages±standard deviation: 42±14 years), 1715 OSA (1,482 males; 54±11 years), 672 insomnia (518 males; 52±14 years), and 649 COMISA subjects (280 males; 56±11 years). Subjects with OSA and COMISA were more obese than the healthy and insomnia groups (p<0.05). On the other hand, insomnia and COMISA groups showed greater depressive mood and greater sleep dissatisfaction (higher PSQI and ISI) compared with healthy and OSA groups. In analysis of PSG parameters, the OSA and COMISA groups showed a higher AHI, a higher ODI, a higher arousal index, a higher percentage of stage 1 sleep and lower percentages of stage 2, 3 and REM sleep than healthy and insomnia groups (Table 2).

**Table 2.** Demographic and clinical characteristics of OSA, insomnia, COMISA and healthy groups

|  | Healthy<sup>a</sup><br>(n=1186) | OSA<sup>b</sup><br>(n=1715) | COMISA<sup>c</sup><br>(n=672) | Insomnia<sup>d</sup><br>(n=649) | p-value<sup>1</sup> | post-hoc<sup>2</sup> |
| --- | --- | --- | --- | --- | --- | --- |
| Group criteria | 20<age<70 | 20<age<70 | 20<age<70 | 20<age<70 |  |  |
|  | ISI ≤ 14 | ISI ≤ 14 | ISI > 14 | ISI > 14 |  |  |
|  | AHI ≤ 15 | AHI > 15 | AHI > 15 | AHI ≤ 15 |  |  |
| No. patients | 1186 | 1715 | 649 | 672 |  |  |
|  | (male: 751) | (male: 1482) | (male: 280) | (male: 518) |  |  |
| **Age** | 43.3 (14.4) | 51.5 (11.3) | 53.5 (11.3) | 48.7 (13.5) | <0.001 | a<d<c<b |
| **BDI** | 11.3 (7.8) | 10.5 (7.2) | 17.7 (9.5) | 18.9 (10.1) | <0.001 | a,b<c<d |
| **AHI** | 6.1 (4.6) | 39.8 (20.6) | 38.6 (22.0) | 6.0 (4.2) | <0.001 | a,d<b,c |
| **ISI** | 7.9 (4.1) | 7.9 (4.0) | 18.5 (3.1) | 19.4 (3.3) | <0.001 | a,b<c<d |
| **Arousal index** | 15.5 (7.0) | 32.8 (16.2) | 33.9 (18.0) | 17.2 (7.1) | <0.001 | a,d<b,c |
| **BMI** | 23.9 (3.4) | 26.7 (3.9) | 26.7 (4.5) | 23.8 (3.6) | <0.001 | a,d<b,c |
| **ODI** | 4.4 (4.1) | 34.1 (20.8) | 32.1 (22.2) | 4.4 (4.0) | <0.001 | a,d<b,c |
| **ESS** | 9.3 (4.7) | 9.7 (4.6) | 11.2 (5.6) | 9.5 (6.0) | <0.001 | a,b,d<c |
| **PSQI** | 6.6 (3.0) | 6.2 (2.6) | 11.1 (3.4) | 12.5 (3.6) | <0.001 | a,b<c<d |
| **N1 %** | 14.1 (7.7) | 26.5 (14.1) | 26.3 (15.1) | 15.3 (7.7) | <0.001 | a,d<b,c |
| **N2 %** | 55.8 (10.0) | 51.0 (12.1) | 51.2 (13.1) | 58.4 (10.6) | <0.001 | b,c<a<d |
| **N3 %** | 8.6 (9.0) | 3.6 (5.5) | 3.7 (6.1) | 5.9 (7.1) | <0.001 | b,c<d<a |
| **REM %** | 21.5 (6.4) | 18.9 (6.6) | 18.9 (7.6) | 20.4 (7.6) | <0.001 | b,c<a,d |

<sup>1</sup> Each p-value was computed for analysis of covariance (ANCOVA) using a general linear model
<sup>2</sup> post-hoc tests were performed between all the pairs of groups. Bonferroni correction was used to adjust p-values considering the number of comparisons.
a, b, c and d in the post-hoc column indicate each group of healthy, OSA, COMISA and insomnia subjects, respectively.

The Institutional Review Board at the Samsung Medical Center approved the retrospective analysis of the deidentified PSG data set without requiring additional consent for its use in this study (IRB No.2021-09-039).

Each sleep EEG data composed of six channels: frontal (F3, F4), central (C3, C4) and occipital (O1, O2). EEG signals are sampled at 200 Hz. We excluded sleep latency intervals from our sleep EEG data. Furthermore, data were fixed at six hours, meaning that data of less than six hours were eliminated, and data of more than six hours were cut off at the six-hour mark. Through these processes, we attempted to minimize waking epoch in the dataset.

### 4.2. EEG preprocessing and artifact removal

We filtered sleep EEG signals using a band pass filter with a range of 0-50Hz and removed artifacts related to ECG and EOG signal components using AAR plugin of EEG LAB.^48^ When the amplitude of signal at each time point was more than five times that of the standard deviation in each channel for the given individual, we considered it as an artifact due to head-motions and it was removed and interpolated. We then performed the z-score normalization by standardizing the amplitude of each EEG channel with respect to the mean and standard deviation of the individual’s whole data. We also excluded the two reference channels from the analysis. Finally, arousal intervals were automatically detected and removed using a previously established arousal detection algorithm that applied a support vector machine classifier to the scalogram built upon EEG data of 16-21Hz frequency range and demonstrated 94-98% of the classification accuracy.^49^

### 4.3. Brain age prediction model (Figure 1A)

#### 4.3.1 Scalogram

Preprocessed EEG data were converted into scalograms based on the continuous wavelet transform method using complex Morlet wavelet function^50^ as shown in Eq. (1)

The complex Morlet wavelet function is given by:

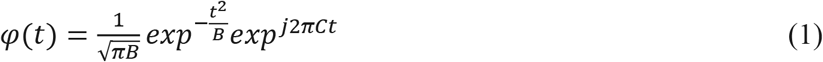

Where *t* is time factor, *C* is the center frequency and *B* is bandwidth.

We chose to use 16 frequency bands of the center frequency *C*, which were determined to be 0.5, 0.7, 0.9, 1.2, 1.6, 2.1, 2.8, 3.8, 5.0, 6.7, 8.9, 11.9, 15.8, 21.1, 28.1, and 37.5 Hz using the log scale distancing. Each band width *B* was determined as 1.5 times of the corresponding center frequency. Finally, we constructed each individual channel’s scalogram using Eq. (1) and the determined *C* and *B*. We down-sampled six hour-long data to 2,160 time bins, with an interval of ten seconds per time bin. This time bin interval, which resulted in the highest model performance, was empirically chosen. Subsequently, our six EEG channels resulted in a final scalogram data in the form of 16 × 2160 × 6, which was used as the input data in 4.2.2.

#### 4.3.2 Brain age prediction model based on DenseNet

To build the brain age prediction model, we employed DenseNet,^51^ which is a CNN architecture using dense connectivity, advantageous in prevention of gradient vanishing, feature propagation enhancement, feature reuse encouragement, and parameter reduction.

The network adopted a previously established and well-tested C-P-DB6-T-DB12-T-DB24-T-DB16-T-AP-FC architecture with 121 three-dimensional layers, in which C is a convolution layer, P is a pooling layer, T is a transition layer, DB is a dense block, AP is an average pooling layer and FC is a fully-connected layer. Each layer is followed by nonlinearity, Rectified Linear Units (ReLU). The model training was performed solely using healthy sleepers (n=1,182). Mean absolute error (MAE) between chronological age and the predicted age was used as the loss function with an Adam optimizer, a learning rate of 0.0001 and a batch size of 2. To train and validate the brain age prediction model, a 10-fold cross validation was used. Source code for the proposed brain age prediction model is available at github link (https://github.com/shyook83/EEG-BAI).

#### 4.3.3 EEG-brain age index

BAI, which reflects a subject’s relative brain health status, was measured by subtracting chronological age from predicted brain age.^22,24,52,53^ Due to regression dilution, however, it is also possible that regression models bias the predicted brain age toward the mean, underestimating the age of older subjects and overestimating the age of younger subjects due to the Gaussian distribution of age in the used dataset. Thus, when deriving the BAI, this bias was corrected using a strategy that was introduced in previous studies.^54,55^

### 4.4 Statistical analysis

#### 4.4.1 Association between electrophysiological brain aging and structural brain aging

To determine whether sleep EEG-BAI is associated with regional cortical thickness, the association was statistically assessed by calculating the BAI and cortical thickness from 49 male participants who underwent PSG and brain MRI on the same day (age [mean±SD]: 42±11 years; range: 21-65 years; demographic information in Table S1). We selected these subjects based on the inclusion/exclusion criteria described in 4.1. We estimated cortical thickness based on the established CIVET pipeline which is a sequential image processing pipeline (CIVET, http://www.bic.mni.mcgill.ca/ServicesSoftware/CIVET)^56^. Briefly, each individual T1-weighted image underwent the following pre-processing steps sequentially: intensity non-uniformity correction,^57,58^ skull-stripping,^58^ registration to the Montreal Neurological Institute (MNI) template space,^59^ and brain tissue segmentation.^53^ Based on the brain tissue labels on image (gray matter, white matter and cerebrospinal fluid), inner and outer cortical surfaces were reconstructed using Constrained Laplacian-Based Automated Segmentation with Proximities algorithm (CLASP).^60^ The inner cortical surface was reconstructed by deforming a spherical mesh onto the white-grey matter (WM/GM) boundary. The outer cortical surface was then reconstructed by expanding the inner surface to the gray matter/cerebrospinal fluid (GM/CSF) boundary and resampled with 40,962 vertices per hemisphere. The cortical surface models were inversely registered into the native space.^61^ We then employed an iterative surface registration to guarantee an optimal vertex correspondence across individuals.^62^ Finally, cortical thickness values were calculated from the Euclidean distance between the inner surface vertices and the corresponding outer surface vertices and subsequently smoothed with a Gaussian kernel with a 30 mm full width at half maximum to improve the signal-to-noise ratio and ensure population analysis.

We then parcellated the cortical surface into seven regions of interest based on human functional networks: sensorimotor, frontoparietal (FPN), dorsal and ventral attention, default mode, salient, language and auditory, and visual network. We then compared a high BAI group (n=14) and a low BAI group (n=35) that were split based on one standard deviation of BAI for the control group (7 years) to analyze the association between BAI and cortical thickness changes in each brain functional region. We used linear models to compare between the two BAI groups while controlling for the covariates (BMI, AHI, chronological age) that were shown to correlate with cortical thickness in the literature.^27,63–66^

#### 4.4.2 Investigation of possible accelerated brain aging in sleep disorders

We performed a linear regression and subsequently Student’s two sample *t*-test to compare BAI between each sleep disorder group and healthy sleeper group. Sex, chronological age and BMI were corrected for group comparison. To investigate the possible association between higher BAI and older subjects in each sleep disorder groups compared to healthy sleepers, we used a linear model and analyzed an interaction term of age × group while effects of sex and BMI were corrected. Furthermore, we sub-grouped each healthy and disorder group using chronological age with a 10-year window and 5-year overlap between the two neighboring subgroups such as 20-30, 25-35, 30-40, 35-45, 40-50, 45-55, 50-60, 55-65, and 60-70. For each age subgroup, we then compared BAI between healthy sleeper and each disorder group to identify the starting point in which sleep EEG aging differs between the groups.

#### 4.4.3 Association of various sleep parameters with sleep EEG-BAI

To quantitatively assess effects of sleep parameters and other covariates (chronological age, sex, depression, BMI) on increased BAI, we conducted a linear regression model. We performed regression analyses between BAI and each parameter/variable for each group separately (e.g., healthy sleeper, OSA, insomnia, COMISA) as well as for all combined groups. The sleep and non-sleep variables analyzed using the linear model included the following: OSA, insomnia, chronological age, sex, BMI, BDI, PSQI, Epworth Sleepiness Scale (ESS),^67^ ISI, percentage of sleep stage (sleep stages 1, 2, 3, REM), and arousal index, as well as the respiratory related sleep disorder indices including AHI and ODI.

#### 4.4.4. Association between the incidence of sleep disorders and EEG-BAI using a hazard model

Kaplan-Meier cumulative incidence curves were used to qualitatively assess the association between BAI and the incidence of OSA, insomnia, and COMISA as chronological age increases. Thus, each hazard model was built based upon the dataset of combining each disorder group with healthy sleeper group. Furthermore, each dataset was divided into four groups (Q1, Q2, Q3, Q4) by sampling an equal number of subjects among Q1-Q4 groups and considering the BAI of individuals. Q1 is the lowest BAI group and Q4 is the highest BAI group. The Cox proportional model was constructed for each disorder to assess association between BAI and sleep disorders while adjusting for chronological age and sex.

#### 4.4.5 Regional spectral power analysis

Our EEG-BAI was observed to significantly increase in OSA, insomnia and COMISA groups compared to healthy sleepers (Results 2.3.). To understand whether the increased EEG-BAI in each sleep disorder is explained by the different pattern of regional EEG spectrum, we analyzed the spatial characteristics of the EEG spectral power among the 6 channels. To compute the spectral power, by applying the 3^rd^ order Butterworth bandpass filter, we differentiated five frequency bands: slow wave (0.5-4 Hz), theta (4-8 Hz), alpha (8-12 Hz), sigma (12-15 Hz) and beta (15-30 Hz). Then, the spectral powers measured for the two channels in each frontal, central, or occipital lobe were averaged. We then compared the spatial characteristics of the spectral power for each sleep disorder group to healthy sleepers by performing a Student t-test independently at each frequency band and each lobe, while adjusting for chronological age and sex.

We also sub-grouped each healthy and disorder group using chronological age with a 20-year window and 10-year overlap between the two neighboring subgroups such as 20-40, 30-50, 40-60, and 50-70. For each age subgroup, we then compared sleep disorder group to healthy sleepers by same method. All the statistical results were corrected for multiple comparisons by controlling the false discovery rate (FDR).

## Data availability

Raw data were generated at the Samsung medical center. Basic demographics and brain age index are available at github link (https://github.com/shyook83/EEG-BAI). Further derived data supporting the findings of this study are available from the corresponding author upon reasonable request.

## Acknowledgements

This work was supported in part by National Institutes of Health (P41EB015922; U01NS086090), Bright Focus Research Grant award (A2019052S) and made possible in part by grant number 2020-225670 from the Chan Zuckerberg Initiative DAF, an advised fund of Silicon Valley Community Foundation and Samsung Medical Center Grant (OTC1190671), the Institute for Basic Science IBS-R029-C3 (NTO1211801) and the Korea Health Technology R&D Project through the Korea Health Industry Development Institute, funded by the Ministry of Health & Welfare, Republic of Korea (HR21C0885).

## Author contributions

S. Yook and H. Kim developed proposed index. S. Yook, H. R. Park, G. Park, E. Y. Joo, and H. Kim performed research. S. Yook, H. R. Park, D. C. Lim, J. Kim, E. Y. Joo, and H. Kim analyzed and validated data. S Yook, H. R. Park, C. Park wrote the paper.

